# *APOE4* genotype negates the benefits of 17β-estradiol on cerebrovascular endothelial and mitochondrial function

**DOI:** 10.1101/2025.07.23.666474

**Authors:** Mackenzie N Kehmeier, Alexandra Famiano, Abigail E Cullen, Thomas Leonhardt, Skylyn Ferguson, Madeleine Snyder, Carrie E McCurdy, Daniel Tyrrell, Nabil J Alkayed, Ashley E Walker

**Author notes:** Contact information: Ashley E. Walker, PhD Department of Human Physiology University of Oregon, 181 Esslinger Hall, 1240 University of Oregon Eugene, Oregon, 97403, USA.

## Abstract

**Background:** Postmenopausal females who carry an *APOEε4* allele are at higher risk of late-onset Alzheimer’s Disease compared to age-matched *APOEε4* males. Estrogen deficiency predisposes females to an increased risk of vascular, cognitive, and metabolic impairments. While estrogen and *APOE* genotype are known to impact metabolic and mitochondrial function in the brain, their cerebrovascular effects are less understood. Thus, the purpose of this study was to determine the interaction between *APOE* genotype and estrogen on cerebrovascular endothelial and mitochondrial function.

**Methods:** Young female homozygous *APOEε3* and *APOEε4* mice (n=19-20/group; ~6 months old) fed a high-fat diet were ovariectomized (OVX), OVX and supplemented with 17β-estradiol, or left intact.

**Results:** In *APOEε3* mice, OVX was associated with impaired posterior cerebral artery endothelium-dependent dilation, which was rescued by 17β-estradiol. However, in *APOEε4* mice, there was no effect of OVX or 17β-estradiol on cerebral artery endothelial function. Carotid artery passive stiffness was greater with OVX and lower with 17β-estradiol treatment in *APOEε3* mice, but there was no impact of OVX or 17β-estradiol in the *APOEε4 mice.* In cerebral arteries and arterioles, mitochondrial complexes I and I+II respiration were lower in *APOEε4* mice compared with *APOEε3* mice. 17β-estradiol led to higher mitochondrial complex I respiration in *APOEε3* but not *APOEε4* mice. These functional differences were concomitant with group differences in mitochondrial DNA copy number, antioxidant enzymes, and pro-inflammatory factors. In contrast to other outcomes, we found that 17β-estradiol treatment was associated with lower cerebral artery stiffness in *APOEε4* but not *APOEε3* mice.

**Conclusions:** Overall, these results indicate that the *APOE* genotype modulates the impact of estrogen on the cerebral vasculature. We found that 17β-estradiol enhances cerebrovascular endothelial and mitochondrial function in *APOEε3* mice but not in *APOEε4* mice. The results suggest that 17β-estradiol supplementation has more cerebrovascular benefit for *APOEε4* non-carriers.

**Novelty & Significance:** *What is known?:*

- Females have twice the risk of Alzheimer’s disease compared with males, and the *APOE4* genetic variant is associated with a greater risk for Alzheimer’s disease compared with the *APOE3* variant.
- The risk for Alzheimer’s disease increases after menopause in females, suggesting that the loss of female sex hormones may play a role.
- There are highly inconsistent results among past studies examining the interaction of *APOE* genotype and estrogens on cognitive function and other brain outcomes.

*What new information does this article contribute?:* Vascular outcomes were not measured in previous studies examining the interaction between *APOE* genotype and estrogens. As such, we aimed to determine the impact of *APOE4* genotype on the cerebrovascular response to estradiol. We found that estradiol improved cerebral artery endothelial function and mitochondrial respiration in *APOE3* mice following ovariectomy. In contrast, *APOE4* mice were refractory to the beneficial effects of estradiol on cerebrovascular endothelial and mitochondrial function. The broader implication of this research is that *APOE* genotype may be a consideration when prescribing hormone replacement therapy to menopausal females due to the impact on vascular outcomes.

## Introduction

*APOE* genotype is the greatest genetic risk factor for late-onset Alzheimer’s disease (LOAD). The *APOE* polymorphic alleles *APOEε2* (E2), *APOEε3* (E3), and *APOEε4* (E4) have a worldwide frequency of 8%, 78%, and 14% respectively(1), and the E4 allele is associated with a 3- to 10-fold greater risk for LOAD(2). Two-thirds of LOAD cases are females, with postmenopausal females carrying an E4 allele having higher rates of LOAD compared with age-matched males with an E4 allele(1), suggesting an interaction between E4 and estrogen deficiency(1, 2). Hormone replacement therapy may serve as a beneficial therapy; however, conflicting reports exist as to how *APOE* genotype modulates the effects of estrogen on LOAD risk. Different previous studies indicate that estrogen has greater effects in E4 than E3(3–5), or the same effect in both E3 & E4(6). Other reports suggest that it has negative effects in E4 while having positive effects in E3(7–9). While the literature is contradictory, studies have not investigated the effects of E4 and estrogen on cerebrovascular dysfunction, a key pathophysiological factor in LOAD.

The first biomarker of LOAD is altered cerebral blood flow, closely followed by metabolic changes in the brain(10). Disruption in blood flow is linked to endothelial dysfunction, resulting from reduced nitric oxide (NO) bioavailability, excessive reactive oxygen species (ROS), and an overall pro-inflammatory milieu(18). As females progress through the menopause transition, endothelial function declines and is restored by treatment with estrogen(13). Estrogen not only enhances NO levels but also supports an antioxidant, anti-inflammatory state and significantly influences mitochondrial function – a primary source of ROS(14). In parallel, metabolic dysfunction is a hallmark of the LOAD brain, stemming from impaired glucose delivery and utilization. This reduced glucose metabolism, detectable in the early stages of LOAD via F-FDG-PET imaging(10, 15), is particularly evident in perimenopausal women carrying the E4 allele, who exhibit a lower metabolic rate than their E3 counterparts(16–18). Additionally, the E4 allele is associated with altered mitochondrial dynamics and dysfunction in astrocytes(19) and neurons(20). Mitochondria influence vascular tone and health(21), possibly linking their dysfunction to the altered cerebral hemodynamics observed in E4 carriers. Old age is associated with impaired cerebrovascular mitochondrial function(22). However, the impact of *APOE* genotype on cerebral vascular mitochondrial function has yet to be investigated. Thus, understanding the interaction between *APOE* genotype and estrogen on the cerebral vascular mitochondria provides novel insight into the vascular dysregulation that occurs with menopause and LOAD.

The purpose of this study was to investigate the interaction effect of *APOE* genotype and estrogen status on cerebral vascular function and metabolism. We hypothesized that E4 female mice would have worse cerebral artery endothelial function (pressure myography), worse cerebrovascular mitochondrial function (Oroboros respirometry), greater large and cerebral artery stiffness (aortic pulse wave velocity [PWV], and passive stiffness), and worse cognitive function (Novel Object Recognition, Morris Water Maze, Nest Building, and accelerating rotarod) than E3 female mice. We further hypothesized that ovariectomy would cause greater deficits in these outcomes in E4 mice compared with E3 mice, and these would be restored in the 17β-estradiol-treated mice. However, in contrast to our hypotheses, we found that vessels from E4 mice do not respond to 17β-estradiol.

## Methods

### Animal Characteristics

We purchased E3 (JAX#029018) and E4 (JAX#027894) mice from the Jackson Laboratories and established breeding colonies. We studied homozygous E3 and E4 female mice at 6 months of age. All mice were on a standard chow diet (Lab Diet, PicoLab Rodent Diet 20, 5053) until they received ovariectomy or sham surgery at 4 months of age, after which, mice were fed a high-fat, high-cholesterol diet (Atherogenic Rodent Diet, Teklad, TD.02028; 42.6% kcal from Fat). Food and water were provided ad libitum, and mice were housed on a 12/12-hour light-dark cycle at 24°C. Mice were euthanized by exsanguination under isoflurane immediately prior to vascular studies. All animal procedures conform with the Guide for the Care and Use of Laboratory Animals (8^th^ edition, revised 2011) and were approved by the Institutional Animal Care and Use Committee at the University of Oregon. See Table 1 in Supplemental Material for animal characteristics, and supplemental methods for more information.

### Ovariectomy & Estradiol Implants

Female mice were randomly assigned to sham (Sham), ovariectomy (OVX), or ovariectomy supplemented with 17β-estradiol (OVX+estradiol). OVX procedures were performed at 4 months of age. One-week post-ovariectomy, a 60-day slow-release 0.36mg 17β-estradiol pellet (Innovative Research of America, Sarasota, FL) was inserted subcutaneously.

### Cerebrovascular Reactivity

Endothelium-dependent vasodilation was assessed *ex vivo* in isolated pressurized posterior cerebral arteries (PCAs), as previously described(23, 24). Arteries were excised and cannulated onto glass micropipettes in a myograph chamber (Danish MyoTechnology Inc.) filled with a physiological salt solution. All the arteries were pre-constricted with phenylephrine (1-6 μM) to obtain 20-40% pre-constriction of luminal diameter. Increases in luminal diameter in response to increasing concentrations of endothelium-dependent dilators acetylcholine (ACh: 1×10^−9^ to 1×10^−4^ M) and insulin (1 to 10000 ng/mL) were determined. ACh and insulin responses were repeated in the presence of Nω-Nitro-L-arginine methyl ester hydrochloride (L-NAME, 30 minutes, 0.1 mM). Endothelium-independent dilation was measured by increases in luminal diameter to sodium nitroprusside (SNP) (1×10^−10^ to 1×10^−4^ M). See Supplemental Table 2 for artery characteristics.

### Cerebrovascular Mitochondrial function

*Ex vivo* cerebrovascular mitochondrial function was measured by high resolution respirometry in O2k instruments (Oroboros, Innsbruck, Austria). Brain arteries and arterioles were carefully dissected from the brain and placed in ice-cold BIOPS (10mM Ca-EGTA buffer, 0.1 µM calcium, 20mM imidazole, 20 mM taurine, 5 mM K-MES, 0.5 DTT, 6.56 mM MgCl2, 5.77 mM ATP, 15 mM phosphocreatine, pH 7.1), followed by permeabilization with 30 ug/mL saponin in BIOPS for 30 minutes, followed with a 30-minute wash in MIRO5. The arteries were then spun down gently and placed in the respirometry chamber. All data were collected at 37° C in a super-oxygenated environment (200-400 µmol/L O2). Mitochondrial integrity was confirmed by measurement of respiratory responses to cytochrome C (inclusion threshold <15%). Each chamber received the same titration protocol probing with carbohydrate-based substrates with the following sequential additions: 5mM pyruvate, 2mM malate, 10 mM glutamate, 5 mM ADP, 10 mM cytochrome c, 10 mM succinate, 1 mM FCCP, 10uM rotenone, 10 μM antimycin A, and 10 μM myxothiazol. This provides an in-depth evaluation of cerebral artery respiratory capacity and substrate control during OXPHOS, as well as the extent of non-phosphorylating respiratory leak, and noncoupled enzymatic capacity of the ETS. Once respiratory measures were completed, the cerebral arteries were removed, spun down and frozen for further analysis.

### DNA Content and mRNA Expression

Cerebral arteries and arterioles from respirometry were used to measure mitochondrial DNA (mtDNA) content and an additional set of cerebral arteries was used to measure mRNA expression, please see the supplemental methods and primer sequences in Supplemental Material Table S3.

### Protein Expression

Protein expression levels were measured for the hippocampus and aorta by Western blot. Detailed methods and antibodies can be found in Supplemental Material.

### Arterial Stiffness

Aortic stiffness was measured *in vivo* by pulse wave velocity (PWV), and detailed methods can be found in the Supplement Material(23). Passive arterial stiffness was measured *ex vivo* in the carotid artery and PCA by changes in lumen diameter and medial wall thickness while increasing intraluminal pressure after a 60-minute incubation in a calcium-free solution(25). From the stress-strain curves, β-stiffness and elastic modulus at low pressure (EM_LP_) and at high pressure (EM_HP_) were calculated as previously described(26).

### Cognitive Function

Spatial memory was assessed via Novel Object Recognition, anxiety via open field, learning via the Morris Water Maze, instinctual behavior via Nest Building, and motor function via accelerating rotarod and velocity during open field and water maze trials. Detailed methods can be found in the supplement.

### Statistical Analysis

Statistical analyses were performed with GraphPad Prism 10. A two-way analysis of variance (ANOVA) was used to determine the interaction effects and independent effects of genotype and estrogen status. In the case of a significant F-value, post-hoc analyses were performed using a Tukey correction for pre-planned comparisons. A repeated-measures ANOVA was used to determine group differences for dose responses. Significance was set at p<0.05, and values are represented as mean±SD. Outliers were identified as z score >2 and were removed from the dataset.

## Results

### Animal Characteristics

E4 mice had a higher body mass than E3 mice (genotype effect p=0.001; Supplemental Table S1). Perigonadal white adipose tissue mass had a main effect of estrogen and genotype (estrogen effect p=0.004, genotype effect p=0.0005) and was highest in the E4 OVX group (Table S1). Confirming successful ovariectomy surgeries, uterus mass and plasma 17β-estradiol were lower in ovariectomized mice (estrogen effect p=0.002, p=0.02). A main effect of estrogen was seen for liver weight (estrogen effect p=0.007), spleen (estrogen effect p=0.0002), and total cholesterol (estrogen effect p=0.04), all higher in the OVX+estradiol group. Plasma progesterone was higher in the Sham groups (estrogen effect p=0.04; Table S1). Lastly, fasting glucose and glucose tolerance did not differ by genotype or estrogen status (Supplemental Figure S1). There was a trend for in vivo oxygen consumption, measured by indirect calorimetry, to be affected by estrogen status (p=0.09) and for energy expenditure to be affected by genotype (p=0.1, Supplemental Figure S1).

### Cerebral artery endothelial function is impaired by ovariectomy in E3 mice but not E4 mice

There was a significant effect of estrogen status on the vasodilatory responses to ACh (p=0.008), but no effect of *APOE* genotype Figure 1A&D). E3 OVX mice had 20% less vasodilation than the E3 sham (max: p=0.005, dose-response: p=0.0002), and 18% less vasodilation than the E3 OVX+estradiol treatment mice (max p=0.002, dose-response: p<0.0001) to ACh. Among the E4 mice, PCA vasodilation to ACh did not differ with OVX or estradiol status (all p>0.05). For the PCA vasodilation in response to insulin, there was a main effect of genotype with the E4 mice having lower vasodilation than the E3 mice (p=0.04); however, there was no interaction and only a trend for estrogen status (p=0.09, (Figure 1B&E). The vasodilation response to ACh and insulin in the presence of LNAME was nearly absent and did not differ between groups (dose-response and maximal responses, all >0.05, Figure 1D&E), suggesting that the group differences in vasodilation are due to differences in NO bioavailability. Endothelium-independent dilation in response to SNP was not different between groups (dose-response and maximal responses, all p>0.05), indicating that the differences in ACh-mediated vasodilation were due to endothelial dysfunction (Figure 1C&F). Sensitivity (EC50) to vasodilators and the amount of pre-constriction also did not differ between groups (all p>0.05, Supplemental Table S2).

**Figure 1.**
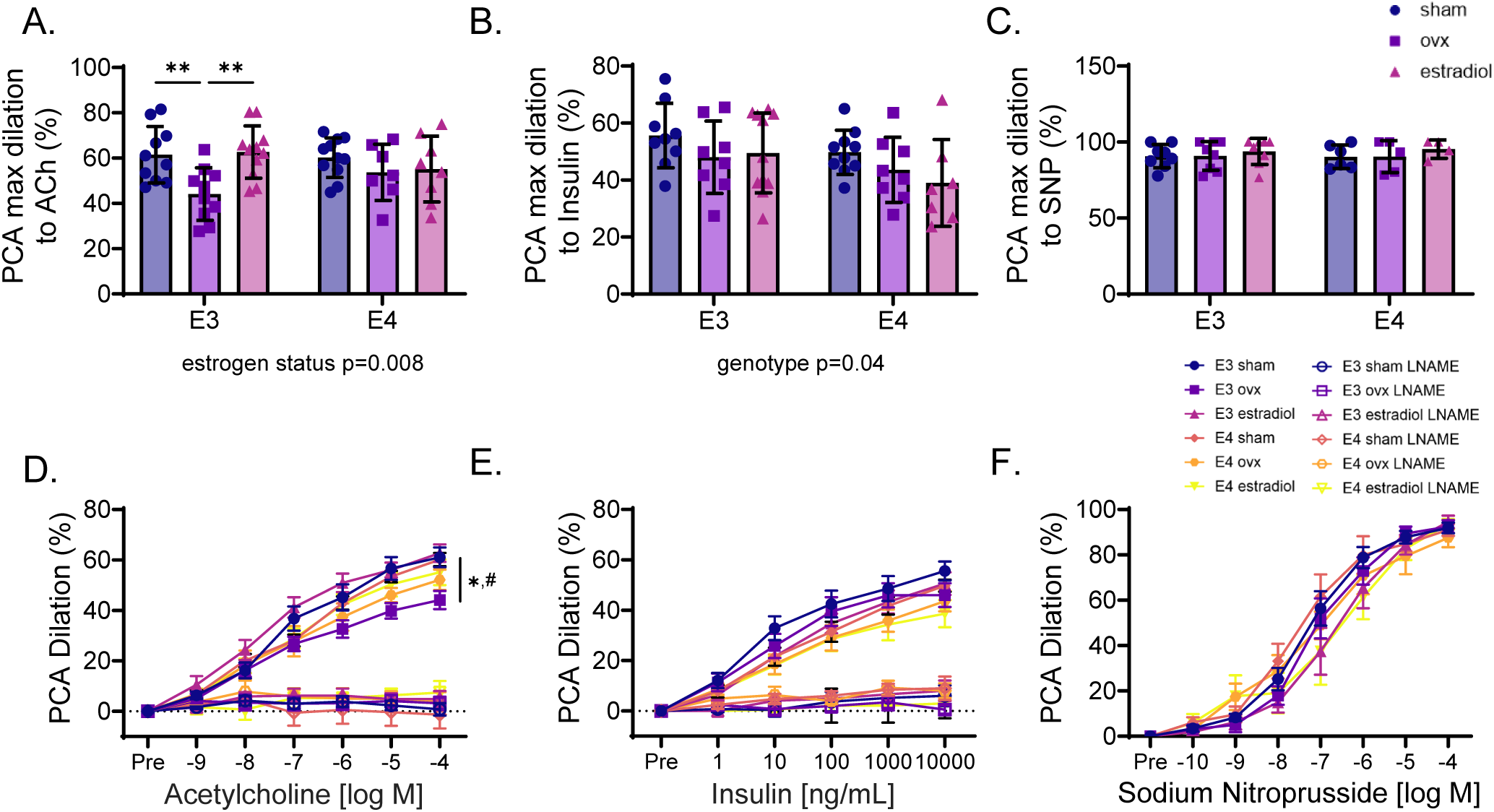
Endothelium-dependent dilation of the posterior cerebral artery is impaired after ovariectomy and rescued by estradiol replacement in *E3* but not *E4* females. Female homozygous E3 and E4 mice were studied at 6mo after receiving a sham, ovariectomy (OVX), or OVX + estradiol. In isolated posterior cerebral arteries (PCAs), endothelium-dependent dilation was measured to A,D) increasing doses of acetylcholine (ACh) and B,E) insulin, in the presence and absence of nitric oxide synthase inhibitor, LNAME. C,F) Endothelium-independent dilation in response to increasing doses to sodium nitroprusside (SNP). A repeated measures ANOVA for the dose responses (* vs E3 sham, # vs E3 OVX+estradiol compared to E3 OVX) and a two-way ANOVA for the maximal responses was used to determine main and interaction effects, and Tukey’s post-hoc analysis was performed. *p<0.05, **p<0.01, ***p<0.001. n=7-11/group. Data are mean±SD.

### Cerebral artery respiration is altered by estrogen status in E3 but not E4 female mice

For the mitochondrial respiration studies, our samples of dissected arteries and arterioles were primarily composed of smooth muscle cells and endothelial cells, as indicated by higher expression of *Acat2* and *Pecam1* in the vascular samples compared with the hippocampus (both p<0.0001). Our vascular samples contained traces of astrocytes (*Gfap*), neurons (*Rbfox3*), and microglia (*A1f1*), but these were significantly lower than the hippocampus (p=0.01, p=0.0008, p=0.005, respectively; Figure 2G).

**Figure 2.**
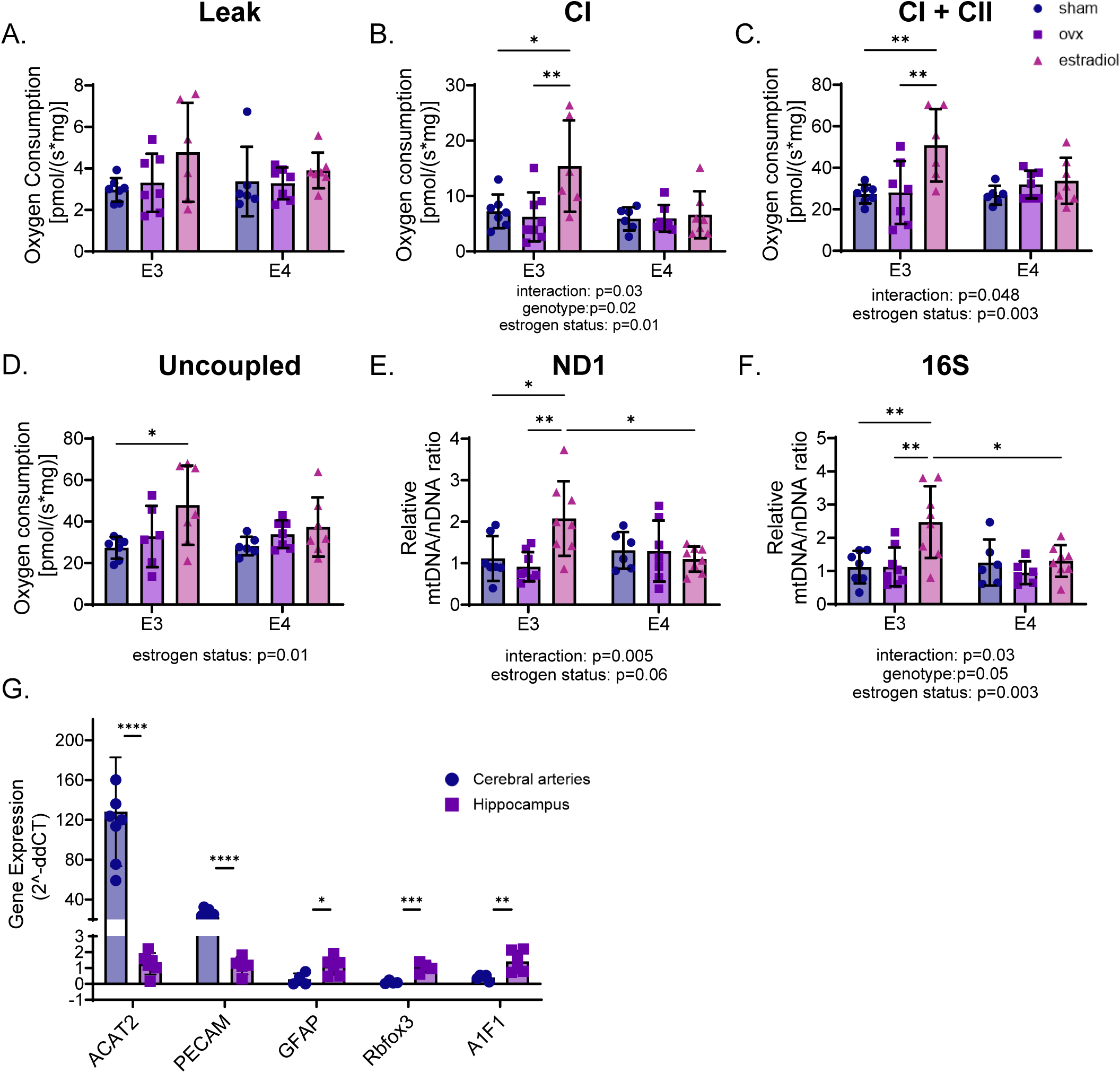
Estrogen status and *APOE* genotype interact to influence cerebral vascular mitochondrial function. Cerebral artery mitochondrial respiration measurements for A) leak, B) complex I (CI) respiration, C) CI+ complex II (CII) respiration, and D) uncoupled maximal respiration after FCCP. Mitochondrial DNA content measured as the ratio to nuclear DNA content for E) 16S and F) ND1. G) Gene expression in cerebral arteries (CAs) and hippocampus homogenates for *Acat2* (smooth muscle cells), *Pecam1* (endothelial cells), *Gfap* (astrocytes), *Rbfox3* (neurons), and *A1f1* (microglia) to demonstrate that respiration samples were enriched for the vascular cells. Gene expression was normalized to hippocampal measures. A one-tailed t-test was used to determine significance between CA and hippocampal samples. A two-way ANOVA was used to determine main and interaction effects in other outcomes, and Tukey’s post-hoc analysis was performed. *p=0.05, **p=0.01. n=5-7/group. Data are mean±SD.

Estrogen status and *APOE* genotype interacted to impact CI coupled respiration (interaction p=0.03) and CI+CII coupled respiration (p=0.05) in cerebral vessels (Figure 2B&C). For CI and CI+CII coupled respiration, there was a main effect of estrogen (CI p=0.01, CI+CII p=0.003), while for CI there was a main effect of genotype (p=0.02, Figure 4.2 B & C). E3 OVX+estradiol mice had greater CI and CI+CII respiration than the E3 sham mice (CI p=0.006, CI+CII p=0.002) and E3 OVX mice (CI p=0.002, CI+CII p=0.002; Figure 2B&C). There was also a main effect of estrogen status in uncoupled respiration, where estradiol supplementation elevated maximal respiration, primarily driven by the E3 OVX+estradiol group (p=0.01; Figure 2D). Interestingly, among E4 mice, CI, CI+CII, or uncoupled respiration did not differ with estrogen status (all p>0.05, Figure 2A-D). There were no differences in leak between groups (all > 0.05, Figure 2A). Following the respiration experiments, mtDNA was measured relative to nuclear DNA in the vascular samples. There was an interaction effect of estrogen status and *APOE* genotype for mtDNA/nDNA content (p=0.03, p=0.005;Figure 2E&F). E3 OVX+estradiol mice had more 16S and ND1 DNA content than E3 shams (p=0.001, p=0.008), E3 OVX (p=0.0007, p=0.0009), and E4 OVX+estradiol mice (p=0.01, p=0.02); however, mtDNA content did not differ in E4 mice with OVX or estradiol.

**Figure 3.**
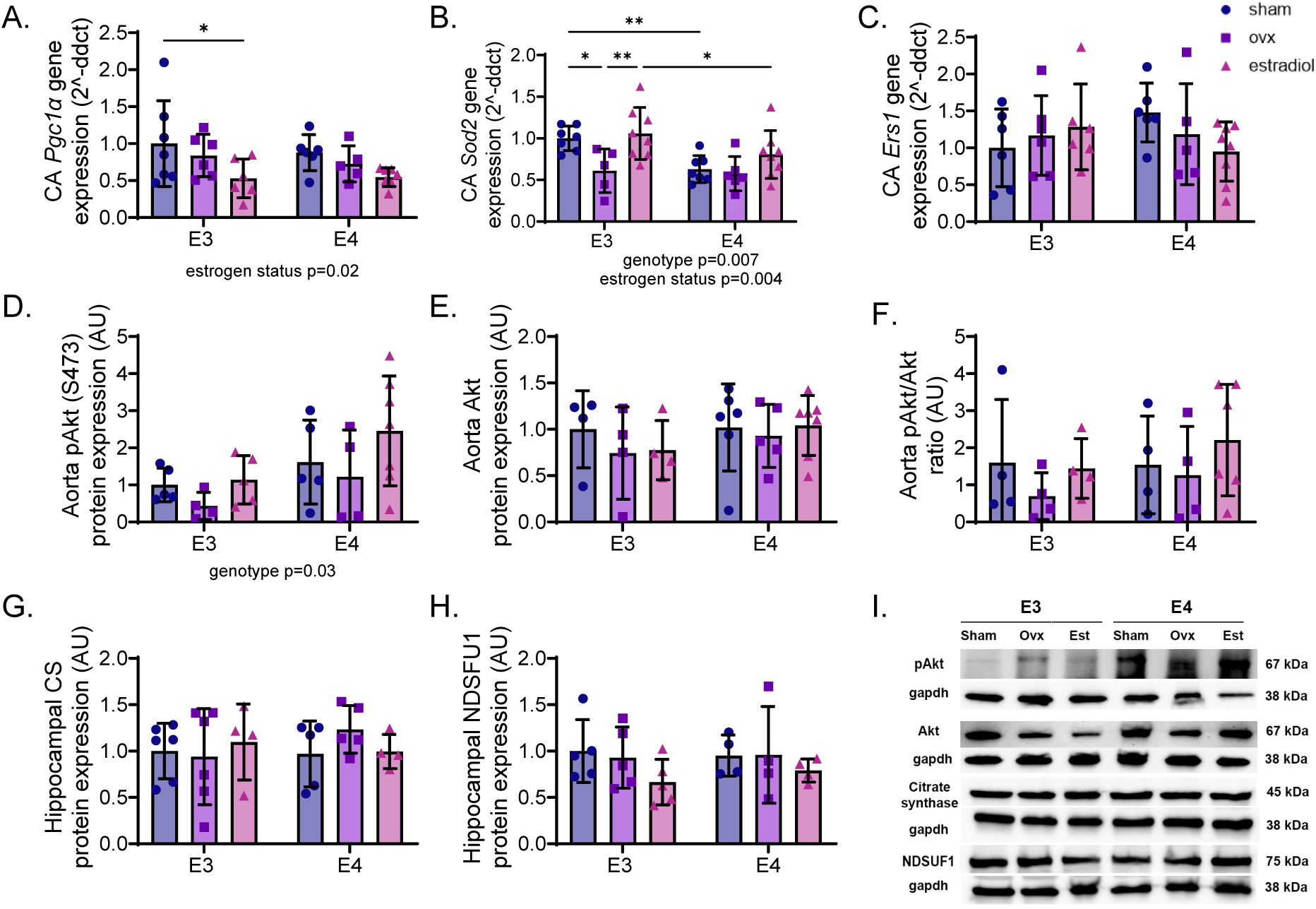
Mitochondrial-associated genes and proteins are altered by *APOE* genotype and estrogen status. Cerebral artery (CA) gene expression for A) Pgc1α, B) Sod2, and C) *Ers1*. Aorta protein expression for D) phosphorylated Akt at serine 473 (pAkt), e) Akt, f) pAkt:Akt ratio. Hippocampal protein expression for G) citrate synthase (CS) and H) mitochondrial complex-I protein NDSFU1. I) Representative blot images. A two-way ANOVA was used to determine main and interaction effects, and Tukey’s post-hoc analysis was performed. *p=0.05. n=3-7/group. Data are mean±SD.

**Figure 4.**
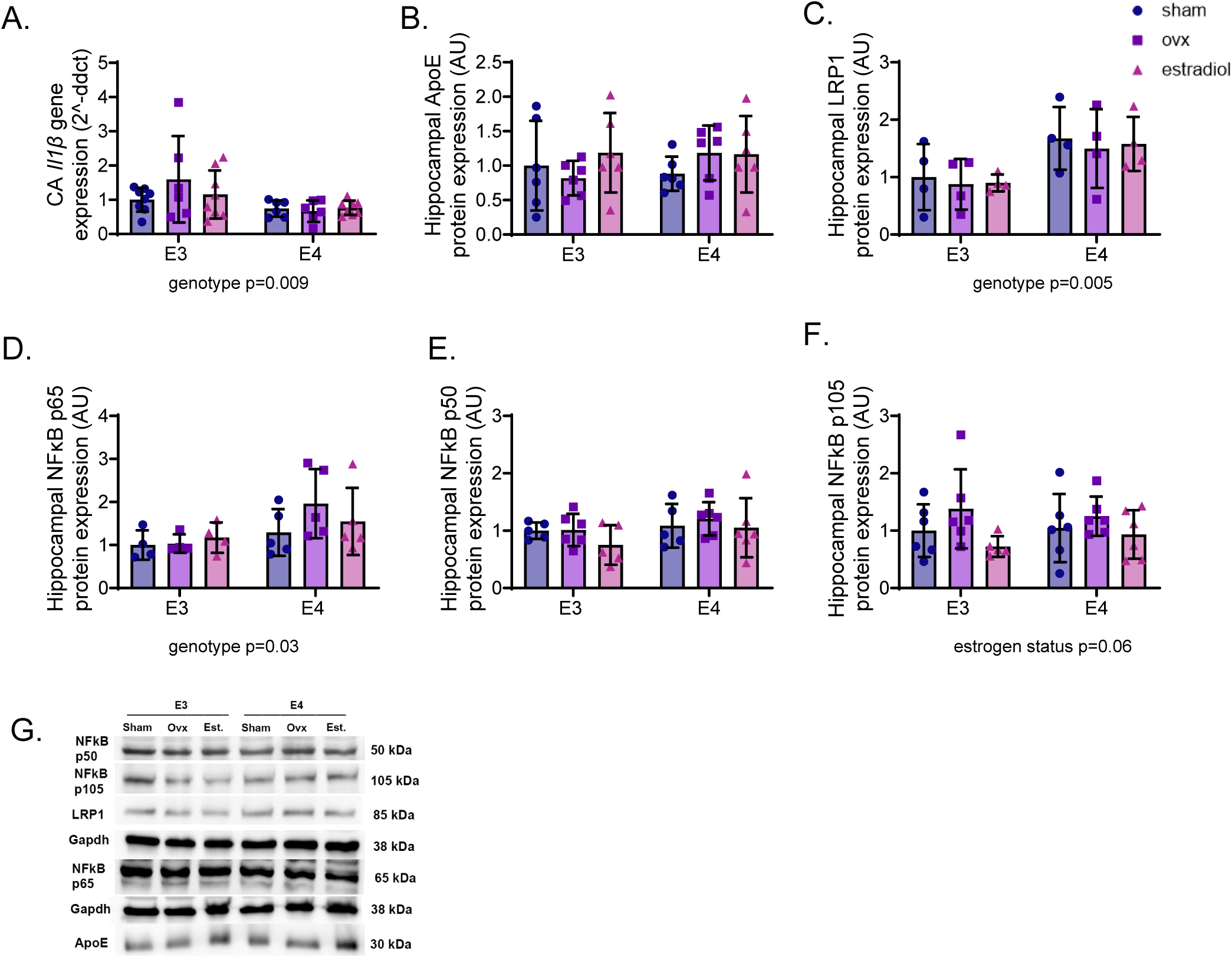
Cerebral artery inflammation is blunted in E4 females, and hippocampal inflammation and ApoE expression are altered by APOE genotype and estrogen status. Cerebral artery (CA) gene expression for A) IL1β. Hippocampal protein expression for B) ApoE, C) LRP1, D) NFkB p65, E) NFkB p50, and F) NFkB p105. G) representative blot images with their gapdh below. ApoE was normalized to total protein. A two-way ANOVA was used to determine main and interaction effects, and Tukey’s post-hoc analysis was performed. *p=0.05. n=3-7/group. Data are mean±SD.

### Mitochondrial factors are impacted by estrogen status and APOE genotype in cerebrovascular tissue

There was a main effect of estrogen status on cerebral artery *Pgc1α*, such that estradiol-treated mice had lower levels (p=0.02), but there were no effects of genotype (Figure 3A). For cerebral artery *Sod2* gene expression, there was a main effect of estrogen (p=0.004) and genotype (p=0.007), such that E3 mice and estradiol-treated mice had high *Sod2* expression (Figure 3B). Specifically, *Sod2* expression was lower in E3 OVX mice compared to E3 sham (p=0.02) and E3 OVX+estradiol mice (p=0.007) but did not differ in E4 mice with OVX or estradiol. Cerebral artery *Ers1* did not differ by *APOE* genotype or estrogen status. Protein expression of aortic phosphorylated Akt at serine 473, had a main effect of *APOE* genotype, being higher in E4 mice compared to *E3* mice (p=0.03), but the total or ratio of Akt did not differ (p>0.05, Figure 3D-F). Protein expression for hippocampal citrate synthase and mitochondrial CI protein NDSFU1 did not differ between groups (p>0.05, Figure 3G&H).

### Cerebral artery and hippocampus inflammatory signaling are altered primarily by APOE genotype

Cerebral artery gene expression of inflammatory cytokine *Il1b* had a main effect of genotype, being higher in E3 mice regardless of estrogen status (p=0.009, Figure 4A). LRP1 protein expression was higher in E4 mice than E3 mice (genotype effect p=0.005, Figure 4C). Total NFκB p65 was higher in E4 mice compared with E3 mice (genotype effect p=0.03, Figure 4D). NFκB p105 expression trended to be influenced by estrogen status, where ovariectomized mice had greater expression (estrogen status effect p=0.06, Figure 4E), but there was no effect of genotype. Protein expression of ApoE and NFκB p50 were neither influenced by genotype nor estrogen status (p>0.05, Figure 4B, 4F).

### Large and cerebral artery stiffness is influenced by estrogen and APOE status

*In vivo* large artery stiffness, measured by aortic PWV, was influenced by the interaction of OVX and *APOE* genotype (p=0.046 Figure 5A); specifically, E4 OVX mice had great aortic PWV compared with E3 OVX mice (p=0.02). We measured carotid artery passive stiffness *ex vivo* and found that β-stiffness did not differ between any groups (all p>0.05, Figure 5B). For carotid artery EM_LP_, there was an interaction effect (p=0.01) and main effect of estrogen status (p=0.01, Figure 5C). E3 OVX mice had a higher carotid artery EM_LP_ than the E3 sham (p=0.001), E3 OVX+estradiol groups (p=0.01), and E4 OVX (p=0.006, Figure 5C). For EM_HP_, no group differences were observed (p>0.05, Figure 5D). These data were generated from stress-strain curves displayed in Figure 5E&F. For PCA passive stiffness, β-stiffness did not differ between groups (all p>0.05; Figure 5G). PCA EM_LP_ and EM_HP_ were dependent on estrogen status and *APOE* genotype (EM_LP_ interaction effect p=0.001; EM_HP_ interaction effect p=0.005; Figure 5H&I). PCA EM_LP_ was greater in E4 OVX mice compared to E4 sham mice (p=0.003) but was not different from the E4 OVX+estradiol mice (p>0.05). These data were generated from stress-strain curves displayed in Figure 5J & K.

**Figure 5.**
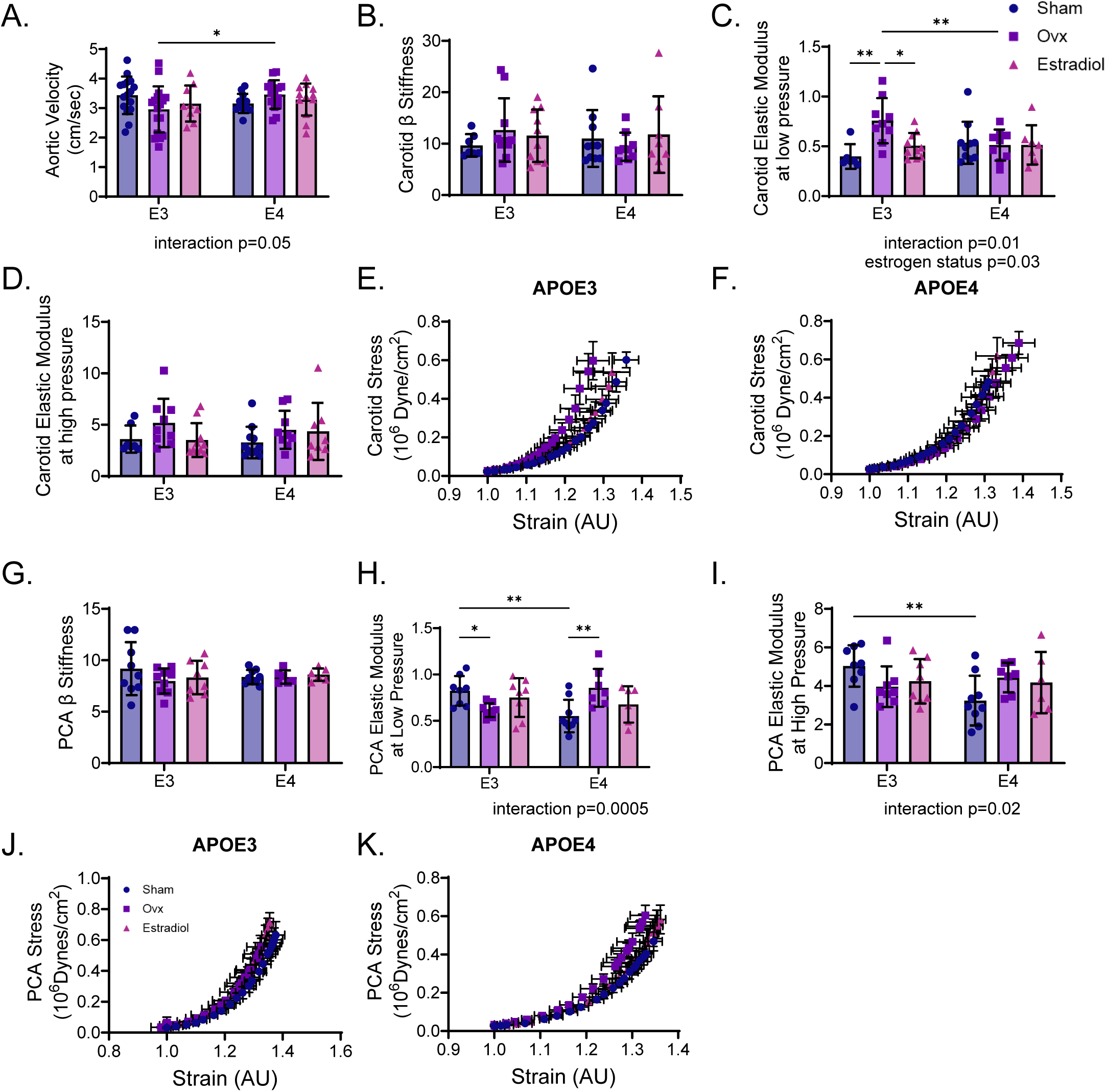
Large artery and posterior cerebral artery stiffness is influenced by *APOE* genotype and estrogen status. A) Aortic stiffness measured by pulse wave velocity (PWV). B) Carotid β stiffness, elastic modulus at C) low pressure and D) higher pressure were calculated from E, F) stress strain curves. Posterior cerebral artery (PCA) G) β stiffness, elastic modulus at H) low pressure and I) high pressure were calculated from J,K) stress strain curves. A two-way ANOVA was used to determine main and interaction effects of estrogen status and genotype on stiffness parameters. *p<0.05, **p<0.01. n=7-18/group, data are mean±SD.

### APOE genotype and estrogen status did not influence cognitive function

Memory, assessed via Novel Object Recognition, was not different between groups (all p>0.05, Figure 6A-C). Anxiety behavior, as indicated by the time spent in the center of the open field arena, was not different between groups (all p>0.05, Figure 6D). Morris water maze test was used to assess learning, no differences were noted between groups in the training trials (all p>0.05, Figure 6F-G). Instinctual behavior, measured by Nest Building, and motor coordination, measured by accelerating rotarod, did not differ between groups (all p>0.05, Supplemental Figure S2). Motor function assessed by the velocity during open field and water maze did not differ between groups (Figure 6E&H).

**Figure 6.**
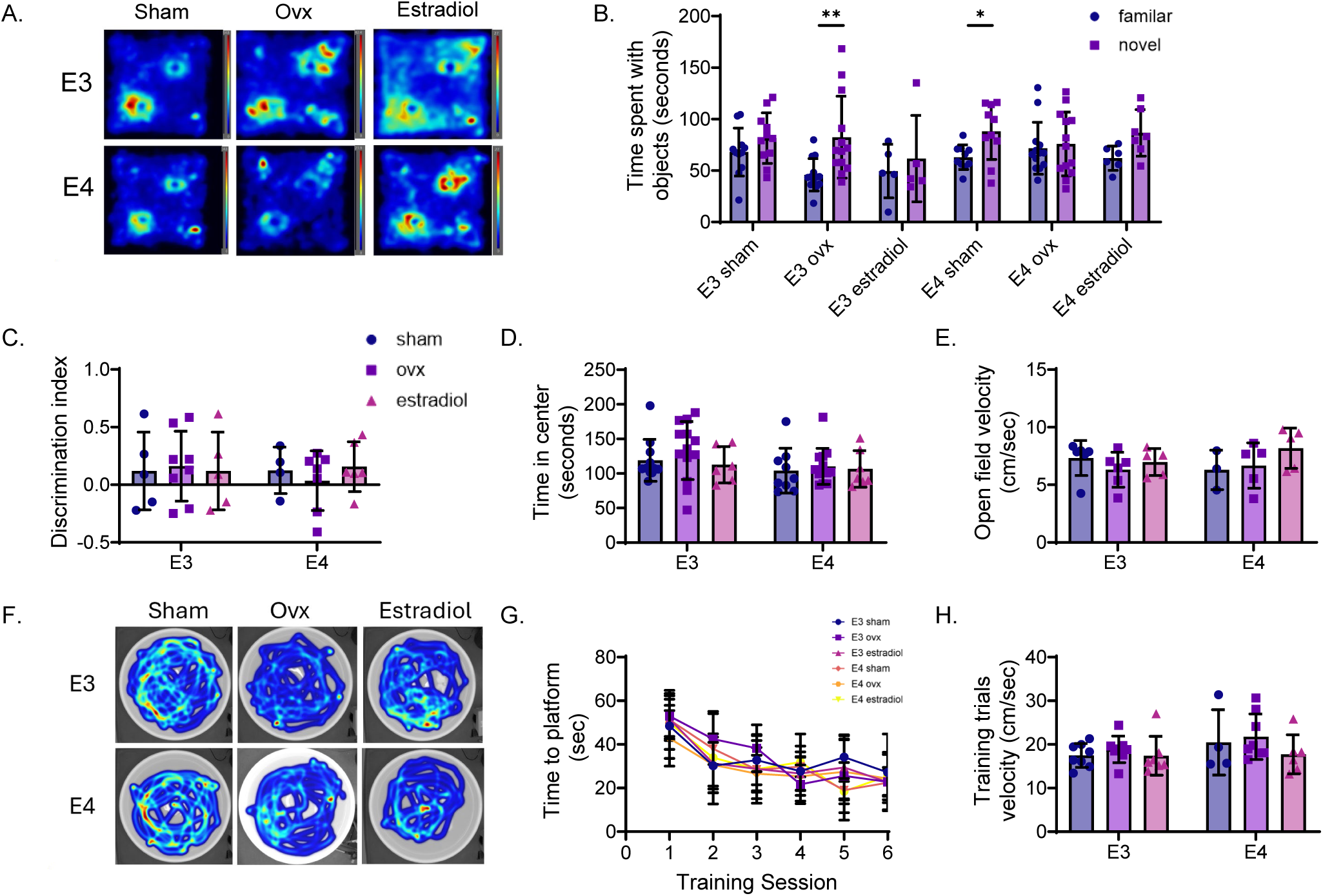
Object memory, anxiety, and learning were not influenced by APOE genotype and estrogen status. Novel object recognition (NOR) A) representative heat maps, B) time spent with familiar (fam) vs novel objects, and C) discrimination index. Open field D) time in center and E) velocity. Morris water maze F) representative heat maps traces for training trials, G) time to platform during training trials, and H) training velocity. A two-way ANOVA was used to determine main and interaction effects. A one-tailed t-test was used for panel (b). *p<0.05, **p<0.01. n=3-8/group, data are mean±SD.

## Discussion

Contrary to our original hypotheses, this study demonstrated that the cerebral vasculature in female mice with the E4 genotype shows resistance to the effects of 17β-estradiol, particularly in relation to endothelial and mitochondrial function. OVX was associated with impaired cerebral artery endothelial function and mitochondrial respiration in E3 mice, and these were rescued by estradiol supplementation. However, E4 mice had no detriments or benefits from OVX or estradiol supplementation in terms of cerebral artery endothelial or mitochondrial function. The impacts of *APOE* genotype and estradiol on endothelial and mitochondrial function may have been driven by differences in mitochondrial abundance, antioxidant defenses, inflammatory signaling, and receptor expression. Estradiol also affected carotid artery stiffness only in E3 mice, and in contrast to other outcomes, estradiol affected cerebral artery stiffness only in E4 mice. *APOE* genotype and estrogen status did not have strong impacts on *in vivo* metabolism, cognitive function, or motor function. In summary, we found that estradiol impacts cerebral artery endothelial function, cerebrovascular mitochondrial function, and large artery passive stiffness in E3, but not E4, female mice.

### E4’s resistance to estrogen: What do we know?

Several previous studies have assessed the effects of *APOE* genotype on the response to estrogen for cognitive function and other neural outcomes. Estrogen replacement therapy improves or maintains memory and brain volume in E4 non-carriers, but these beneficial effects are not seen in E4 carriers(27–29). In some cases, estrogen replacement therapy results in worse cognitive function and a decline in brain volume in E4 carriers compared with non-carriers(30–32). Similar patterns are found in preclinical models. In E3 mice, supplementation with estradiol is associated with better memory, greater hippocampal neuron spine density, higher brain-derived neurotrophic factor, fewer Aβ plaques, and less microglia-induced inflammation; however, these beneficial effects of estradiol were not seen in E4 mice,(7, 8, 33–36) suggesting that the E4 mice are resistant to the effects of estradiol. In some cases, estradiol had adverse effects in the E4 mice, such as more Aβ plaques after estradiol treatment(8). However, it should be noted that there is some inconsistency in the literature, with a few studies finding that estrogen replacement therapy improves memory and processing speed, maintains brain volume, and reduces Aβ deposition in human E4 carriers(3, 4, 37) and E4 mice(5, 35, 38). Other studies find no influence of *APOE* genotype on cognitive outcomes(39–42). These inconsistencies may be due to differences in age, diet, or disease stage in humans or the presence of Aβ in mice. Despite the wealth of studies examining the interactions of *APOE* and estrogens, no previous studies examined vascular outcomes.

### Endothelial Function

Our data suggest that estradiol supports NO-mediated vasodilation in young E3 female mice but has no effect in E4 mice. However, it is important to note that this pattern was found solely for ACh-mediated vasodilation, and not vasodilation to insulin. In this study, estradiol did not rescue insulin-mediated vasodilation for either genotype, while we previously found that a high soy diet preserves cerebral artery dilation to insulin after ovariectomy in wildtype C57BL/6 mice(43). The mice in the present study were fed a high-fat diet, while in the previous study, we used a normal chow (low-fat) diet, suggesting that a high-fat diet may impact the influence of OVX and estradiol on insulin responses. Overall, our findings suggest that cerebral arteries from E4 mice are resistant to the impacts of estradiol on endothelial function.

### The impact of cerebrovascular mitochondrial function on cerebral vessel dynamics

Mitochondrial health impacts vascular function by regulating the energy source for vasodilation and vasoconstriction, and as a primary producer of ROS(22). This is particularly important in the cerebral vasculature, as brain endothelial cells contain more mitochondria than endothelial cells in other vascular beds(44). Cerebral artery and arteriole respirometry by high-resolution respirometry has only been performed once previously, to our knowledge(22), highlighting the novelty of this technique. Our findings suggest that estrogen supplementation in E3 mice increases mitochondrial CI respiration but does not impact respiration in E4 mice. Mitochondrial CI+CII respiration was also higher in E3 estradiol-treated mice, which appears to be driven by the increase in CI respiration rather than CII respiration. We also find that estradiol supplementation in E3 mice resulted in higher mitochondrial DNA content, suggesting either more or larger mitochondria, a finding not observed in E4 mice. Somewhat counterintuitive, we observed lower *Pgc1α* gene expression in the cerebral arteries from the estradiol-treated groups; however, this could suggest negative feedback due to the higher mitochondrial abundance. Mitochondrial fission and fusion dynamics are potential mediators of our mitochondrial abundance findings, and future investigations are needed to determine these dynamics in the cerebral vasculature of *APOE* and estradiol-supplemented mice.

Previous studies have examined the effects of estrogen and APOE genotypes, separately, on mitochondrial proteins. We found no differences in mitochondrial CI protein NDSFU1 expression between groups, potentially because this protein is transcribed from nuclear DNA, not mitochondrial DNA. In contrast to our findings, a previous study found that mtDNA-encoded CI in cerebral blood vessels was elevated by estradiol treatment in rats (45). Another study found that nDNA-encoded CI protein NDUFB8 was more highly expressed in E4 compared with E3 brain endothelial cells (46). We also measured citrate synthase expression in the hippocampus, the rate-limiting enzyme of the citric acid cycle, and found it was not modulated by estrogen or APOE genotype. Potential explanations for our lack of findings for mitochondrial-related proteins may be due to differences in mitochondrial content of the hippocampus compared to the cerebral arteries. Due to our limitations in the size of the cerebral arteries, protein abundance studies were conducted in the hippocampus, whereas functional studies were performed on the cerebral arteries. Thus, while we did not find effects of estradiol or *APOE* genotype on mitochondrial-related proteins in hippocampus homogenates, it is possible that *APOE* genotype and estrogen have a specific impact on the vasculature.

The mitochondria, primarily CI and CIII, are the primary sites of ROS production. Even though estradiol supplementation increases CI respiration in E3 mice, this does not necessarily result in greater oxidative stress, as estrogen also increases antioxidant enzyme expression(47). Mitochondrial associated SOD (*Sod2*) has previously been observed to be increased by estrogen supplementation in mice(47), similar to our finding in cerebral arteries of E3 mice. We also observed that the E4 genotype is associated with lower *Sod2* expression, suggesting a potential lower resilience to oxidative stress in cerebral arteries with the E4 genotype.

### Other mechanisms that contribute to vascular function

*APOE* genotype is known to influence the expression of receptors and inflammatory mediators(48). Our findings indicated no difference in apoE expression in the hippocampus, which is contrary to past studies demonstrating lower apoE expression in E4 compared with E3(49, 50). ApoE’s primary receptor in the brain is LRP1. We found that LRP1 is elevated in the E4 females and not influenced by estrogen status. The E4 variant of apoE binds less efficiently to LRP1 than the other variants, and perhaps this lower binding affinity triggered the increase in expression of LRP1. ApoE/LRP1 signaling leads to NFκB inhibition, and the reduced efficiency of E4 ApoE binding to LRP1 potentially explains elevations in NFκB in E4 mice in our study. While our results for NFκB in the hippocampus align with previous findings, our findings of lower expression of pro-inflammatory cytokine *Il1β* in cerebral arteries from E4 mice are contradictory. Our findings suggest that the impact of *APOE* genotype on inflammatory status of the cerebral blood vessels may differ from that of the surrounding brain tissue.

### Arterial Stiffness

Increases in arterial stiffness are a potential mediator of cerebrovascular dysfunction and cognitive decline(51). We found interactions of *APOE* genotype and estradiol on vascular stiffness. We observed a higher passive stiffness in the carotid artery following ovariectomy that was rescued by estradiol treatment in the E3 group but not in the E4 group. This aligns with a recent study, where ovariectomy increased carotid passive incremental modulus in OVX mice(52). These changes to carotid artery stiffness could influence cerebral artery endothelial function, as recent studies suggest that age-related changes to carotid compliance have a greater influence on cerebrovascular function than changes to aortic stiffness in females(53). In the cerebral arteries, we found the opposite, where the E4 groups were impacted by estrogen status, but not the E3 groups. Aortic stiffness measured by PWV also appeared to be more affected by OVX in E4 mice, although there were no differences with estradiol treatment for this outcome. Thus, artery stiffness is the only outcome where we find that the E4 mice were more sensitive to changes in estrogen status; and further investigations are needed to understand the mechanisms underlying these effects.

### Cognitive and motor function

While we found no differences in cognitive function between groups in this study, previous studies have found that E4 mice have cognitive deficits in multiple behavioral and learning tasks(54). When examining the impact of APOE genotype and estrogen on cognitive function, a previous study found that estrogen was associated with better performance on a Novel Object Placement test in E4 mice, but not when mice were on a high-fat diet, and estrogen had no impact on E3 mice(55). Thus, the high-fat diet fed to the mice in our studies may explain our lack of findings.

## Limitations

There are some limitations to our study that should be noted. The first limitation of our study is regarding the normalization of the respirometry outcomes by tissue mass. The mass of the cerebral vessels was measured as accurately as possible, but given their small size, there is likely some error. As such, we also normalized these outcomes to nDNA and leak and found similar patterns of group differences regardless of the normalization protocol (data not shown). Second, we did not measure testosterone or aromatase; therefore, we cannot determine the impact of these on our outcomes. Since adipose tissue can produce estrogen, and the body mass of E4+OVX mice was significantly greater than that of E3 mice, there is a possibility that adipose-derived estrogen confounded our results. However, this was not reflected in circulating 17-β estradiol concentrations or uterus mass. Third, mice lack cholesterol ester transfer protein (CETP). CETP exchanges cholesterol and triglycerides between lipoproteins, and the absence of this protein in mice results in them carrying the majority of their lipid cargo in HDL rather than atherogenic LDL particles. Thus, future studies investigating *APOE* genotype should be conducted in the presence of CETP to increase the translation to humans.

## Conclusions

Overall, our results indicate that *APOE* genotype modulates the impact of estrogen on the cerebrovasculature. We found that 17β-estradiol enhances cerebrovascular function and mitochondrial function in E3 mice but not E4 mice. The results suggest that estradiol supplementation may provide greater therapeutic benefit for E4 non-carriers. Further research is needed to understand the mechanisms by which E4 is resistant to estradiol supplementation, which will improve our understanding of disease progression and aid in the development of more effective therapeutics for LOAD.

